# The Mutational Dynamics of Short Tandem Repeats in Large, Multigenerational Families

**DOI:** 10.1101/2021.11.22.469627

**Authors:** Cody J. Steely, W. Scott Watkins, Lisa Baird, Lynn B. Jorde

## Abstract

Short tandem repeats (STRs) are tandemly repeated sequences of 1-6 bp motifs. STRs compose approximately 3% of the genome, and mutations at STR loci have been linked to dozens of human diseases including amyotrophic lateral sclerosis, Friedreich ataxia, Huntington disease, and fragile X syndrome. Improving our understanding of these mutations would increase our knowledge of the mutational dynamics of the genome and may uncover additional loci that contribute to disease. Here, to estimate the genome-wide pattern of mutations at STR loci, we analyzed blood-derived whole-genome sequencing data for 544 individuals from 29 three-generation CEPH pedigrees. These pedigrees contain both sets of grandparents, the parents, and an average of 9 grandchildren per family. Using HipSTR we identified *de novo* STR mutations in the 2^nd^ generation of these pedigrees. Analyzing ~1.6 million STR loci, we estimate the empircal *de novo* STR mutation rate to be 5.24*10^−5^ mutations per locus per generation. We find that perfect repeats mutate ~2x more often than imperfect repeats. *De novo* STRs are significantly enriched in *Alu* elements (p < 2.2e-16). Approximately 30% of STR mutations occur within *Alu* elements, which compose only ~11% of the genome, and ~10% are found in LINE-1 insertions, which compose ~17% of the genome. Phasing these *de novo* mutations to the parent of origin shows that parental transmission biases vary among families. We estimate the average number of *de novo* genome-wide STR mutations per individual to be ~85, which is similar to the average number of observed *de novo* single nucleotide variants.

## Introduction

Short tandem repeats (STRs), or microsatellites, are 1-6 base pair (bp) motifs of repeating units of DNA. These loci make up approximately 3% of the human genome (Lander et al. 2001). STRs are distributed throughout the genome and are located in both coding and non-coding regions (Subramanian et al. 2003). STRs have recently been associated with gene expression, where length variation can regulate gene expression of nearby loci (Gymrek et al. 2016; Quilez et al. 2016; Fotsing et al. 2019). STR expansions are also known to contribute to a number of diseases, including amyotrophic lateral sclerosis (DeJesus-Hernandez et al. 2011; Renton et al. 2011), Huntington disease (MacDonald et al. 1993), fragile X syndrome (Fu et al. 1991), and nearly 50 others (reviewed in (Usdin 2008; Nelson et al. 2013; Hannan 2018)).

STRs have been shown to have high mutation rates when compared to other types of variants, including single nucleotide variants (SNVs) and indels (Legendre et al. 2007; Eckert and Hile 2009; Lynch 2010). The mutation rate for STRs can vary significantly depending on the motif length at the locus of interest (Chakraborty et al. 1997). Their high heterozygosity has made STRs a valuable tool in forensics. Typically, only 13 loci are needed to uniquely identify any person (Jobling and Gill 2004). Multiple mechanisms have been proposed to explain this high mutation rate, including unequal crossing over in meiosis, retrotransposition-mediated mechanisms, and strand-slippage during replication (reviewed in (Fan and Chu 2007)). It is possible that each of these mechanisms contributes to the high mutation rate of STRs, but strand slippage is the mechanism proposed for generating most observed mutations in STR loci (Schlötterer and Tautz 1992). Generally, studies of STR mutation rates have analyzed a small number of loci (Weber and Wong 1993) or have focused on loci on the Y chromosome (Heyer et al. 1997; Zhivotovsky et al. 2004; Ballantyne et al. 2010; Willems et al. 2016). While recent work has examined genome-wide STR mutations in a small number of individuals (Willems et al. 2017) or in disease cohorts (Trost et al. 2020; Mitra et al. 2021), further analysis of STRs is needed to better understand their mutational dynamics in the genomes of healthy individuals.

Due to the repetitive structure of STRs and their high mutability, sequencing and genotyping these loci is difficult, especially using short-read sequencing data. Many tools have been created during the last decade to genotype and identify mutations at STRs and longer tandem repeats across the genome (Gymrek et al. 2012; Dolzhenko et al. 2017; Willems et al. 2017; Dashnow et al. 2018; Mousavi et al. 2019). Some of these tools are designed to detect STR expansions at disease-related loci, while others detect expansions and contractions of STRs genome-wide but are constrained by sequencing read length and the STR motif size.

The three-generation structure of the Centre d’Etude du Polymorphisme Humain (CEPH) pedigrees has been valuable for previous work on mutation rates of single nucleotide variants, mobile element insertions, and structural variants (Feusier et al. 2019; Sasani et al. 2019; Abel et al. 2020; Belyeu et al. 2021). These data have also been used in analyses of the role of maternal age and DNA damage in generating germline mutations (Gao et al. 2019) and in examining the association between SNV mutation rate and longevity (Cawthon et al. 2020). Here, we present pedigree-based empirical estimates of the rate of mutation, parent-of-origin transmission differences, interfamilial repeat length variation, and the distribution of new STR alleles for microsatellite loci throughout the genome using these well-characterized CEPH families.

## Results

We utilized whole-genome sequencing data from 544 individuals in 29 CEPH pedigrees. These pedigrees include three generations, generally with both sets of grandparents in the first generation, the parents in the second generation, and the grandchildren in the third generation (Figure 1). The average number of grandchildren in the third generation is approximately nine (ranging from 7 to 16). We analyzed *de novo* STR mutations in the second generation of these families, and the large number of individuals in the third generation allowed us to analyze and verify transmission of putative *de novo* mutations.

**Figure 1.**
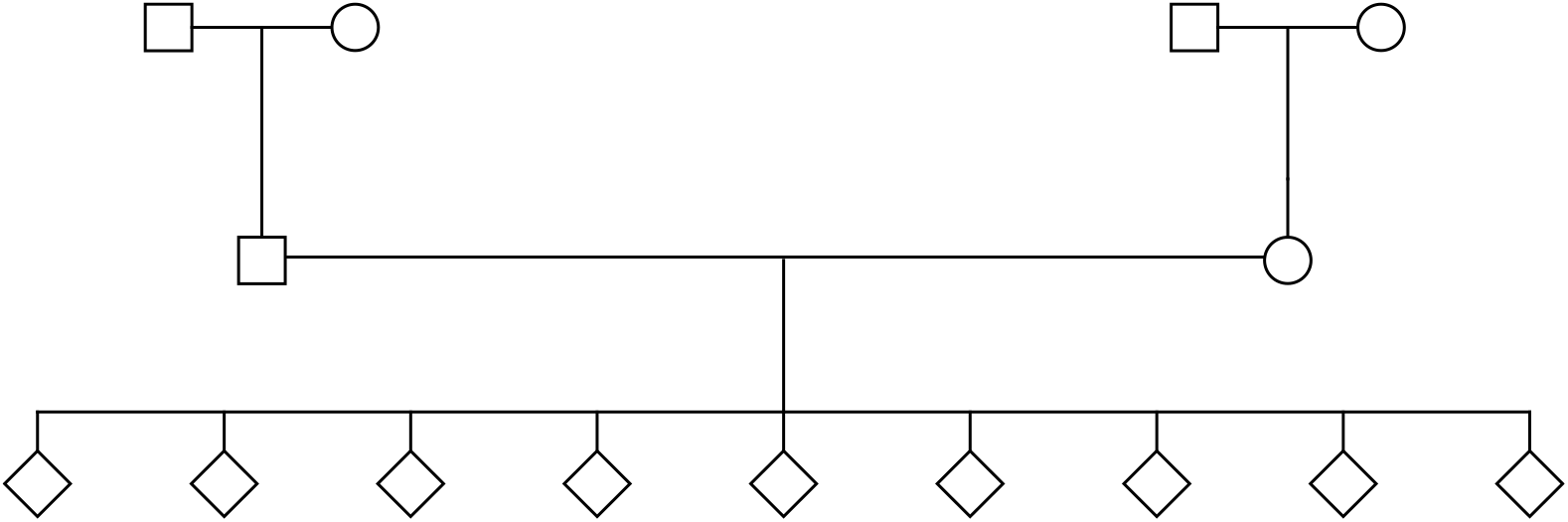
Example CEPH pedigree. Most CEPH pedigrees include both sets of grandparents, both parents, and an average of ~9 grandchildren. This example pedigree has been anonymized and shows the average number of grandchildren in the third generation.

We used HipSTR to genotype and analyze STR loci throughout the genome. HipSTR genotypes each STR and provides the precise length of each allele, but it does not attempt to genotype STRs that are longer than the length of the sequencing read. The precise estimation of STR length is important for analyzing *de novo* mutations that may differ by a single repeat unit. We used a reference file containing more than 1.6 million defined STR loci (see Methods), each of which HipSTR attempted to genotype. We were only able to assay STRs that were present in the reference file. While it is likely that there are other unannotated STRs, the number of STRs in the HipSTR reference file exceeds the number of STRs presently annotated in the human genome reference sequence (hg19). On average, ~49% of STR loci passed our filtering criteria for members of the second generation (see Methods) and could be examined for *de novo* mutation events. Loci that did not pass our filtering generally had low coverage, low posterior probability supporting the genotype, high level of PCR stutter, or a large number of flanking indels.

To assess the accuracy of the genotypes produced by HipSTR, we compared a subset of the genotypes to previously analyzed PCR-based genotypes in the CEPH families (see Methods). We compared the PCR-based genotypes at five random loci in three families (175 total genotypes) to the genotypes generated by HipSTR. The filtered HipSTR genotypes matched 171 of the 175 previously generated STR genotypes for a concordance rate of 97.71%. These previously genotyped loci do not include mononucleotide repeats, which are more difficult to sequence and validate.

In the second generation of each family, we identified *de novo* mutations at STR loci and then traced the transmission of the mutation to the third generation to ensure that it was a germline mutation. We analyzed 68 individuals in the second generation of 29 families (some large families have more than two individuals in the second generation). Those who were excluded were missing a parent in the first generation or had a parent who could not be analyzed by HipSTR (see Methods). Collectively, we were able to identify 5,249 putative *de novo* mutationsin these individuals. We filtered these mutations to ensure that they were transmitted to at least two individuals in the third generation and filtered loci where the parent (in generation 2) without the *de novo* mutation was missing a genotype or shared the same genotype. Approximately 20% of identified putative *de novo* mutations were not found in more than one individual in the third generation, ~22% of the total *de novo* mutations were shared with the other parent in the second generation, and for ~2% of these mutations, the other parent in the second generation had missing data. After filtering these loci, we identified 2,859 *de novo* STR alleles in these individuals for an average of ~42 mutations per individual (Figure 2A). There was a large amount of variation among individuals, with fewer than 10 mutations identified in some individuals and others having nearly 100; however, the number of new mutations discovered per individual follows a normal distribution (Shapiro-Wilk test, p= 0.08). The mutation rates calculated for STR loci show a similar pattern (Figure 2B). The genome-wide mutation rates for STR loci that passed our filtering (which varied by trio) ranged from 5.58*10^−6^ to 1.2*10^−4^ with an average value of 5.24*10^−5^ mutations per locus per generation.

**Figure 2.**
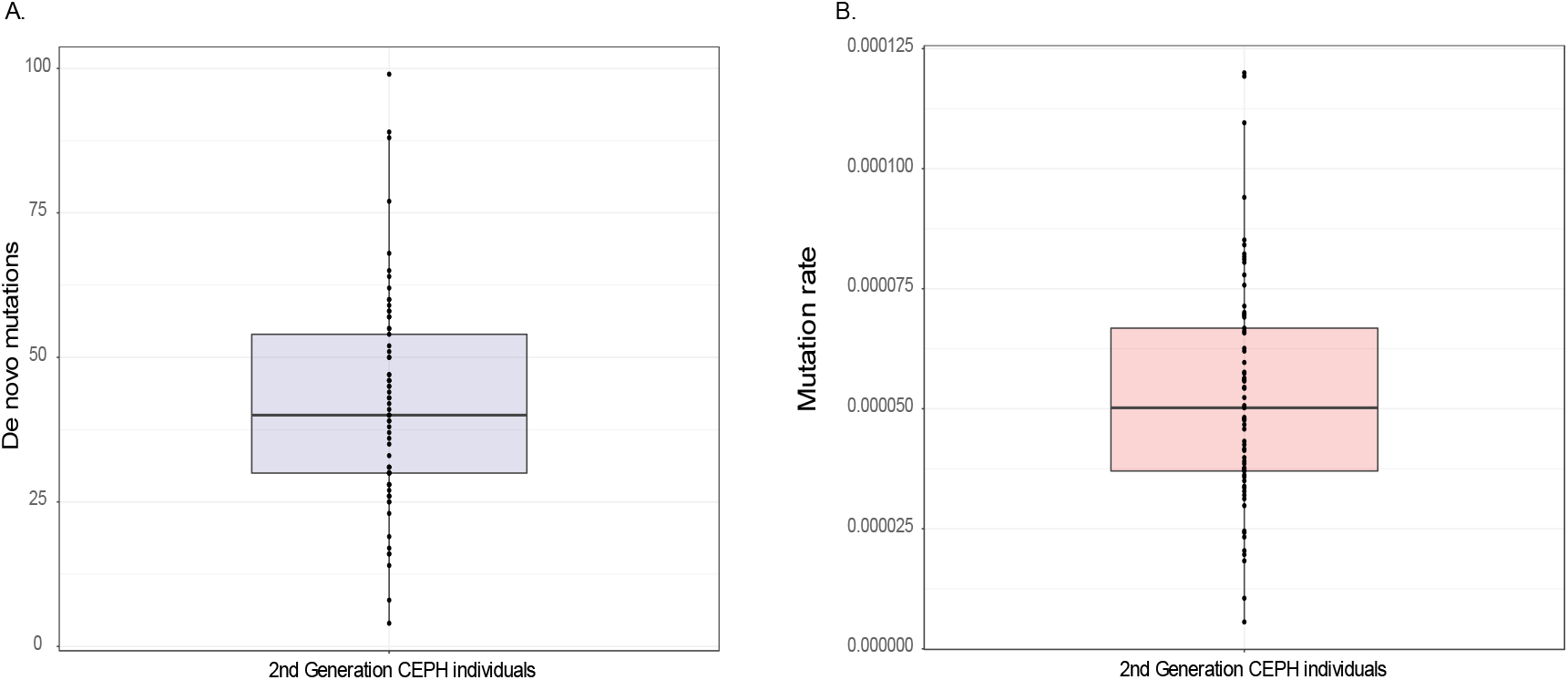
Number of *de novo* mutations and mutation rates among 68 individuals. A) The number of *de novo* STR mutations identified in each individual from the second generation of the CEPH pedigrees. The mean number of identified mutations was ~42, with wide variation in the number of de novo mutations detected per individual. B) The STR mutation rate for each individual in the second generation of the CEPH pedigrees.

Next, we analyzed all unique *de novo* STR mutations passing all filters by their motif length. We divided the number of *de novo* mutations for each motif length by the number of STRs that passed our filters for that length. We found that STRs with shorter motif lengths generally had higher mutation rates than those with longer motif lengths (Figure 3). The mutation rates ranged from 9.99*10^−6^ for hexanucleotide repeats to 7.88*10^−5^ for dinucleotide repeats. Mononucleotide repeats had a slightly lower mutation rate than dinucleotide repeats at 6.82*10^−5^, but a smaller proportion of these passed our filtering methods.

**Figure 3.**
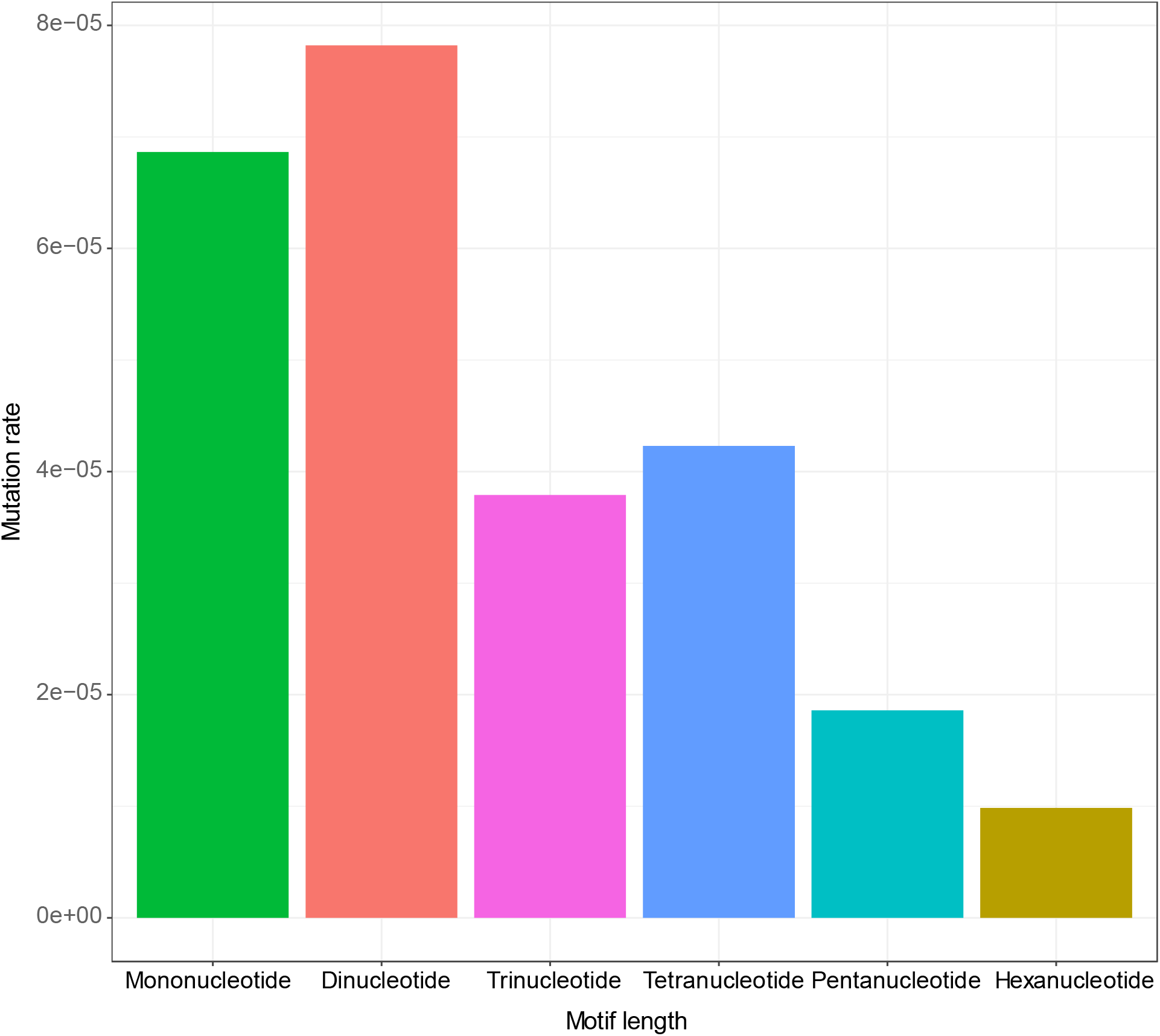
STR mutation rates for different motif lengths. Mutation rates for 2,747 unique genotyped short tandem repeats with motif lengths from 1 bp (mononucleotide) – 6 bp (hexanucleotide). Mutation rates generally decrease as motif length increases, with some exceptions.

We also compared the identified *de novo* STRs that were perfect repeats (e.g. “ATATAT”) against imperfect repeats, those with an interrupted repeat structure (e.g. “ATACAT”). Of the 2,747 unique STR loci with a *de novo* mutation, 2,045 (~74.4%) were classified as perfect repeats, and 702 (~25.5%) were imperfect repeats by Tandem Repeats Finder (Table 1). We found that perfect repeats (0.00213 *de novo* mutations/ total perfect repeat loci) were approximately twice as likely to mutate as imperfect repeats (0.00106 *de novo* mutations/ total imperfect repeat loci) (Two-proportions Z-test, p < 0.00001).

**Table 1.** Perfect and imperfect *de novo* STRs. The number of *de novo* STR mutations that were defined as perfect and imperfect in our dataset. The “Total perfect” and “Total imperfect” columns show the number of genome-wide perfect and imperfect repeats identified. The fractions of perfect and imperfect loci with a *de novo* mutation were calculated by dividing the number of *de novo* mutations by the total number of perfect or imperfect repeats.

|  | Perfect<br>*de novo* | Imperfect<br>*de novo* | Total<br>perfect | Total<br>imperfect | Fraction of perfect<br>loci with *de novo*<br>mutation | Fraction of imperfect<br>loci with *de novo*<br>mutation |
| --- | --- | --- | --- | --- | --- | --- |
| Mono | 788 | 145 | 615604 | 216599 | 0.0013 | 0.00067 |
| Di | 770 | 230 | 143570 | 153117 | 0.0054 | 0.0015 |
| Tri | 132 | 38 | 40510 | 39788 | 0.0033 | 0.00096 |
| Tet | 287 | 230 | 102591 | 135956 | 0.0028 | 0.0017 |
| Penta | 56 | 36 | 37281 | 61933 | 0.0015 | 0.00058 |
| Hexa | 12 | 23 | 20956 | 52125 | 0.00057 | 0.00044 |

After identifying the *de novo* STR mutations in CEPH individuals, we examined the location of these mutations in the genome. Using the UCSC Genome Browser, we identified all exons, introns, 3’-UTRs, and 5’-UTRs in the genome (hg19). We then intersected the *de novo* STR’s position with each of these locations (Figure 4). The majority (53.38%) were found in intergenic regions. Slightly less than half (44.87%) were found in intronic regions, with a much smaller portion being found in UTRs. Only two mutations were found in exons: a trinucleotide (CCG) repeat mutation in *USP24* and a second trinucleotide (CGG) repeat mutation in *PHLPP1*. We compared the ratio of the number of observed and expected *de novo* events found in each of the five genomic features shown in Figure 4. *De novo* STR events are significantly underrepresented in the coding regions of exons (p < 1e-6) but not in the 5’- and 3’-UTRs. In non-coding regions, slightly more *de novo* events were found in introns than expected by chance (Supplemental Table 1).

**Figure 4.**
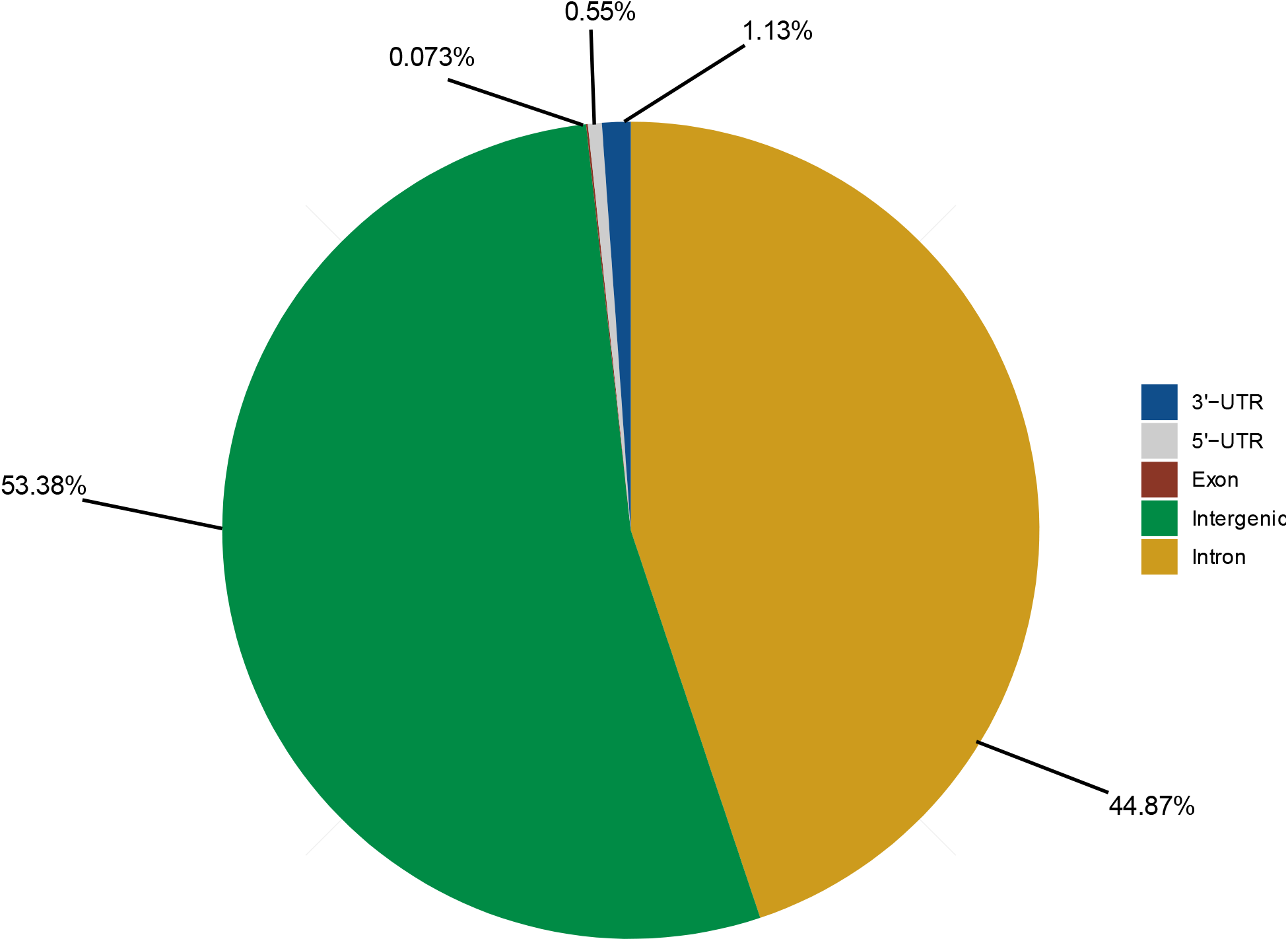
Location of *de novo* STR mutations in the genome. The majority of *de novo* STR mutations were identified in intergenic regions (shown in green) and intronic regions (shown in yellow). Only two mutations overlap with exons.

We also intersected the *de novo* STR locations with transposable element insertion locations in hg19 to determine how often transposable elements (*Alu* elements, LINE-1, and SVA) were potentially the source for new mutations (Supplemental Figure 1). These three families of transposable elements were selected for analysis because they are active in humans. Approximately 30% of the *de novo* STR mutations were found in recognized *Alu* elements throughout the genome, with a smaller fraction of these mutations in LINE-1 (6%) and SVA (0.12%) insertions. We compared the ratio of the number of observed and expected *de novo* events found in *Alu* elements (which compose 11% of the genome), LINE-1 (which compose 17% of the genome), and SVA (which compose 0.1% of the genome). *De novo* STR events were significantly overrepresented in *Alu* elements (p < 2.2e-16) and significantly underrepresented in LINE-1 insertions (p < 2.2e-16). The number of *de novo* events identified in SVA insertions was not significantly different from the expected value (p = 1) (Supplemental Table 2).

To determine if *de novo* STR mutations are more likely to originate in males or females, we analyzed the sex of the grandparent transmitting the most probable haplotype with the *de novo* mutation event (see Methods). Of 2202 resolved haplotypes, 1117 de novo STR alleles were transmitted by males and 1085 were transmitted by females. This slight male transmission bias was not statistically significant (male/female ratio = 1.03, p-value ≥ 0.5, two-sided binomial test).

Individual families, however, varied in their male/female transmission ratios, and 61% of families had elevated male/female transmission ratios (Figure 5). Comparing the mothers and fathers within each family also showed that only four had a statistically significant transmission bias. Families 1421 and 8819_8820 showed female transmission bias, and families 8095_8097 and 8100_8101 showed male transmission bias (p < 0.05, Bonferroni corrected). In three of these four families, a two- to three-fold higher rate of *de novo* transmission in one grandparent accounted for the elevated ratio. This result suggests that some individuals transmit new STR mutations at an atypically high rate. We also analyzed the relationship between parental age and STR mutation rate but did not find a strong correlation for either paternal age (r^2^ = 0.044) or maternal age (r^2^ = 0.0071).

**Figure 5:**
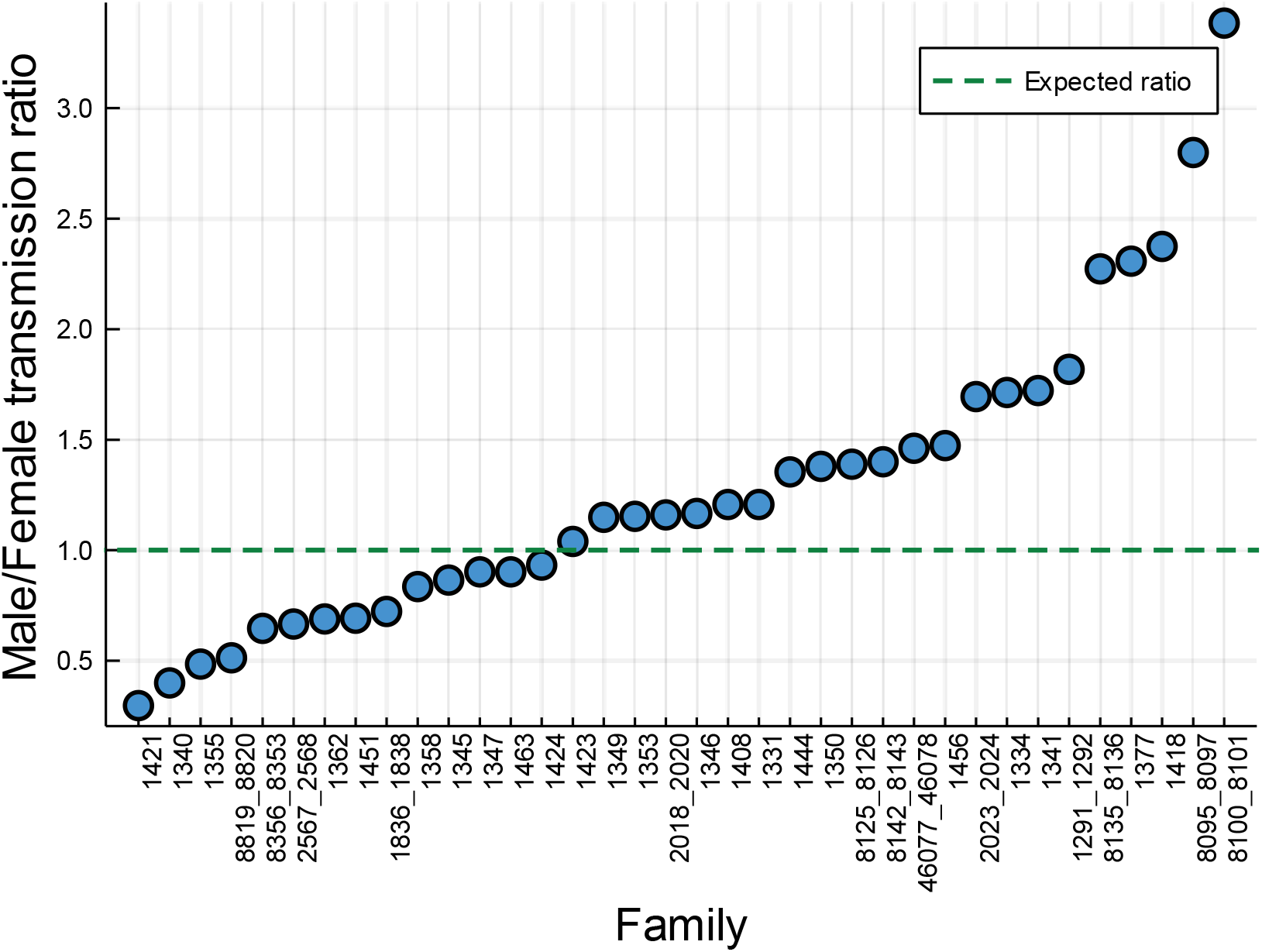
Observed male/female de novo STR transmission ratio by family. The ratio of male to female transmissions of de novo STR alleles is shown by family. The male/female transmission ratio ranged from 0.30 to 3.4. Of the 36 individual families within the 29 large CEPH pedigrees*, 22/36 (61%) have a higher-than-expected rate of de novo STR transmission by males, while 14/36 (39%) have a low-than-expected transmission rate. The male/female transmission ratios suggests a trend for a male transmission bias in the CEPH families. *Several of the 29 CEPH families have an extended pedigree structure which were separated into 36 pedigrees as shown in Figure 1 (see also Methods).

We analyzed the spectrum of size differences between the original and the *de novo* STR alleles for 1388 mutations in which the transmitting haplotype and the size change in base pairs could be confidently identified. Except for three-base pair repeats, smaller mutations were generally more frequent than larger mutations, consistent with the overall pattern of observed mutations (Figure 6a, b). There was not a significant difference between the length of de novo alleles in males and females (p-value ≥ 0.7, two-tailed t-test). Expansions were favored slightly over contractions, primarily due to an excess of 1 and 2-bp expansions (Figure 6c). The stepwise changes observed for all new STR alleles follow a negative exponential distribution. We fitted the number of observed de novo alleles to the size change in base pairs, *f(x)* = *1095.97(exp(−0.521796)(x)*, where x is the de novo allele size change and *f(x)* is the observed number of alleles. The observed number of de novo alleles for all size classes falls within the estimated 90 percent confidence interval. All classes except for trinucleotide changes fall within the 95 percent confidence interval (Supplemental Figure 2). Collectively, new STR alleles rarely differed by more than 6 bp from their original length, consistent with strand-slippage by a single repeat unit across all classes of microsatellites.

**Figure 6:**
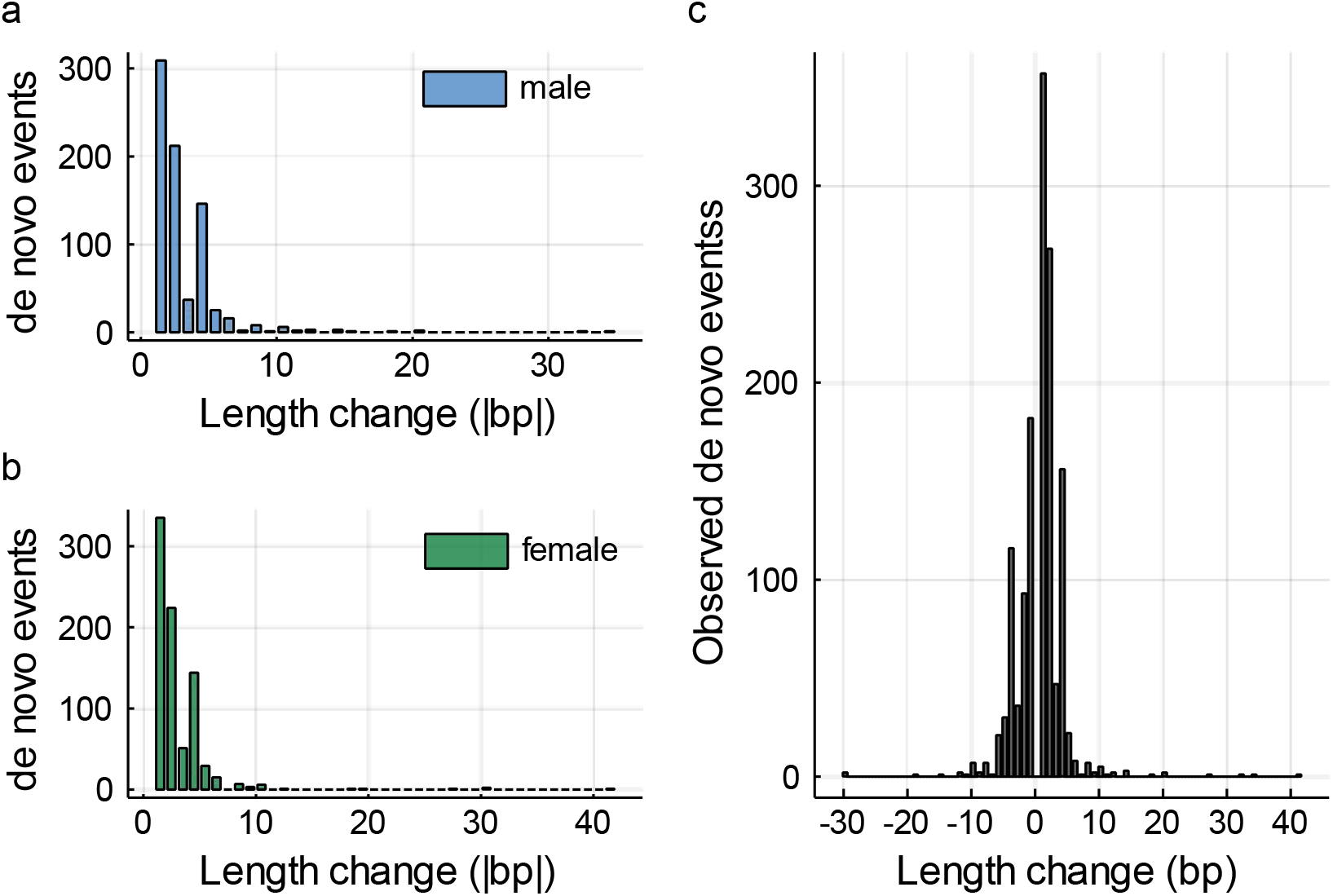
Distribution of length changes for phased *de novo* STRs. a,b) The distribution of length differences between new and original phased STR alleles is similar among CEPH males and females. c) In general, contractions and expansions decrease in frequency with increasing allele size.

After determining the parent of origin for the mutations, we analyzed the change in size, as measured by the number of repeats, between the transmitting grandparent’s allele and the *de novo* allele in the parent explicitly by repeat motif size (Figure 7). The majority of the identified mutations show a single stepwise change, defined as a mutation resulting in a change of a single repeat motif (that is, a dinucleotide repeat expanding or contracting by two bases or a trinucleotide repeat expanding or contracting by three bases, etc.). This pattern is seen for each motif size examined, with the total number of events decreasing as motif size increases.

**Figure 7.**
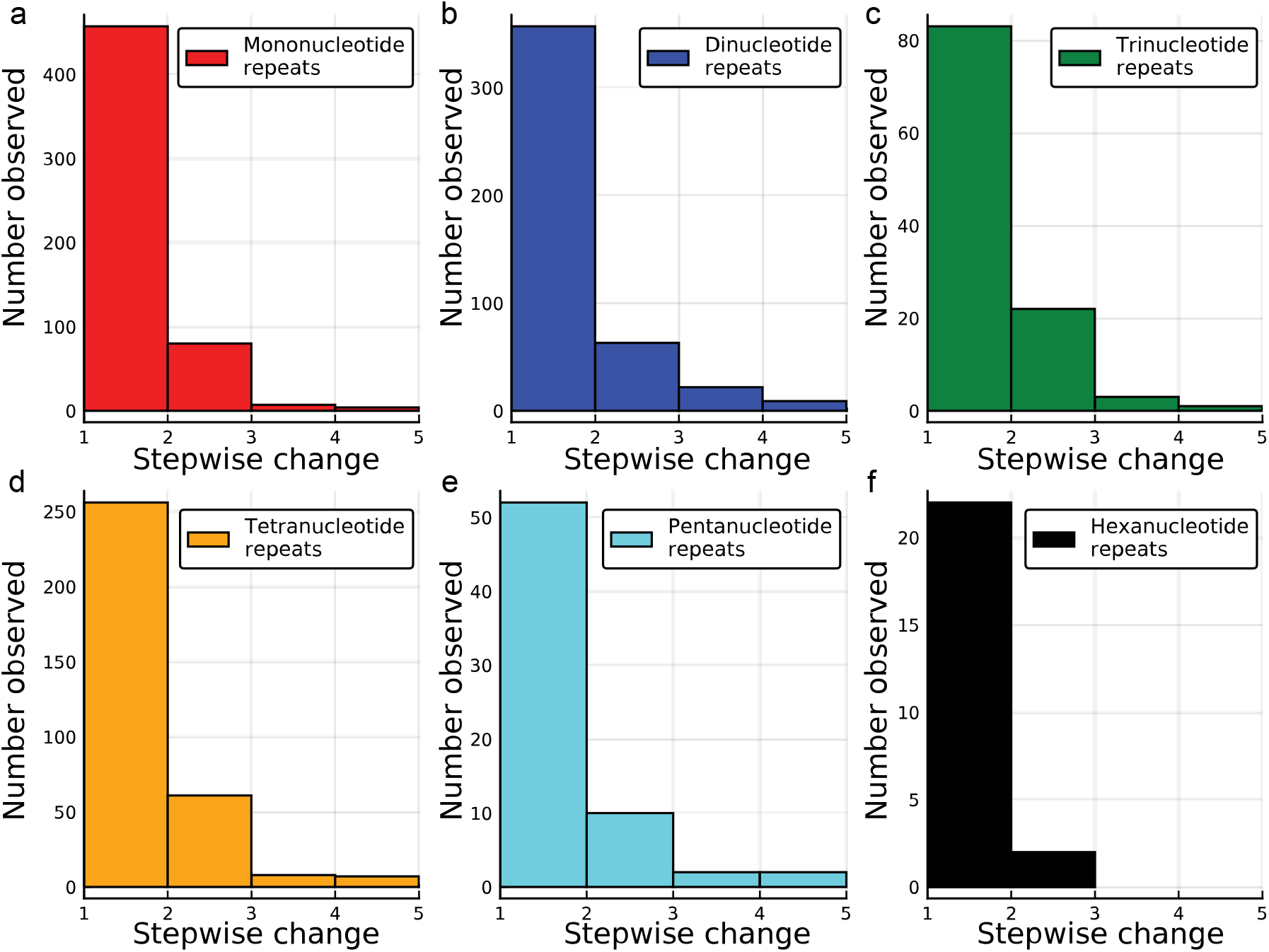
The distribution of stepwise changes for phase known de novo STR alleles by repeat motif size. Panels (a-f) show the distributions of transmitted phased-determined de novo STR alleles for each STR motif length in the CEPH families. In each case, the majority of the observed size changes are a single repeat unit. For all motif-size categories, the frequency of observed de novo STR alleles decreases exponentially with an increasing number of repeats.

## Discussion

We used the unique structure of the CEPH pedigrees to determine the genome-wide mutation rate and the number of STR mutations inherited in 68 individuals from 29 families. Our analysis reveals a high degree of variability among families in the number of transmission-verified *de novo* STR mutations (Figure 2A) and in the mutation rate of STR loci (see Figure 2B). This variation is similar to the pattern seen for single nucleotide variants in these families (Sasani et al. 2019); it may be due to individual differences in genetic backgrounds or differences in DNA repair efficacy. We compared the number of *de novo* STRs to the number of *de novo* single nucleotide variants transmitted by each individual and found no correlation between the number of mutations found (r^2^ = 4*10^−5^). We may not see a relationship between these values due to the small sample size included in our study, or differing DNA repair mechanisms involved in these two mutation types.

Analyzing approximately 49% of all STR loci in the genome, we found an average of ~42 *de novo* STRs in the examined individuals. With our average mutation rate of 5.24*10^−5^ per locus per generation, if we were able to assay all STRs across the genome, we estimate there to be an average of approximately 85 *de novo* STR mutations per individual. This estimate of *de novo* STR mutations in these individuals falls within the range of previous genome-wide estimates (Willems et al. 2016; Willems et al. 2017). However, our estimate is likely to be conservative. This is due in part to the limitations of HipSTR, as we are only able to confidently sample those repeats that are smaller than the sequencing read length (~150 bp), allow for flanking sequence to be mapped, and have reads that completely span the repeat. We also removed alleles that were shared with the other parent or were not passed down to multiple grandchildren in the third generation. Some of the identified mutations that were passed down to fewer than two grandchildren may have been true *de novo* mutations or the product of mosaicism. While it is likely that some true *de novo* STRs were excluded because of this filtering criterion, this prevented a number of false positives from being included in the dataset. Additionally, few loci on the Y chromosome passed our filtering critera, possibly due to the highly repetitive nature of the Y chromosome (Tilford et al. 2001; Skaletsky et al. 2003). While these filtering steps increased confidence in our genotyping, they also decreased sensitivity.

Although many studies have examined the mutation rates of different STR motifs, there appears to be no consensus on how mutation rate and motif size are related. Some studies show that dinucleotide repeats mutate more quickly than the longer tetranucleotide repeats (Chakraborty et al. 1997; Kruglyak et al. 1998), while others show that tetranucleotide repeats mutate more quickly (Weber and Wong 1993; Sun et al. 2012). In our dataset, the mutation rate generally decreases with increased motif length (Figure 3); however, there are exceptions to this pattern. Over all loci, the mononucleotide repeats examined in this study appear to mutate more slowly than the dinucleotide repeats. This is likely due to an under-sampling of the mononucleotide loci, and sampling of only those loci that are smaller than read length. Given the low complexity of these regions, they are more difficult to sequence and confidently genotype. Many of these loci did not pass the filtering applied due to low quality scores or a small number of reads spanning the repeat. Other mononucleotide repeats are located in the tail of transposable elements and are difficult to map accurately, likely contributing to the decreased number of repeats passing our filters. The second STR motif that does not follow this pattern is trinucleotide repeats, which have a slightly lower mutation rate than tetranucleotides. The trinucleotide mutation rate falls within the 90% CI of the best-fit curve for *de novo* allele size changes using all phased *de novo* mutations (Supplemental Figure 2). This finding indicates that mutational steps of three base pairs are not exceptionally low compared to expectation. Trinucleotide repeats are enriched in coding regions relative to mono-, di-, tetra-, and pentanucleotide repeats (Subramanian et al. 2003) and may be more highly conserved due to their genomic location.

The majority of *de novo* STR mutations were found in intergenic or intronic regions, with a very small proportion of these events occurring in exons and 3’- or 5’-UTRs (Figure 4). Only two of these mutations occur in exons, and unsurprisingly, they are both trinucleotide repeats. One of these mutations was found in *PHLPP1*, which has been associated with colorectal cancer (Liu et al. 2009), and the other was found in *USP24*, which has been associated with Parkinson disease (Li et al. 2006). These loci did not pass our filtering in most families, but appear to be polymorphic within the families that did pass our filtering criteria.

Examining non-exonic STRs, we found that many of the identified *de novo* STR mutations overlap with transposable elements (TEs) (Supplemental Figure 1). TEs have been proposed to act as seeds for microsatellites (Arcot et al. 1995; Jurka and Pethiyagoda 1995) because of their poly(A) tails; furthermore, *Alu* elements have an A-rich region in the middle of the element. Approximately 30% of our *de novo* STRs overlap with *Alu* elements, likely in the poly(A) tail, so this may be an underappreciated source of origin for STR loci. This finding supports recent work showing that many non-reference or rare tandem repeat loci are in close proximity to *Alu* elements (Fazal et al. 2020). We found fewer *de novo* STRs in L1s and SVAs, despite the fact that L1s compose a greater portion of the genome than *Alu* elements (reviewed in (Xing et al. 2013)). It is unclear if this difference is due to a lower number of copies of L1 throughout the genome, thus creating fewer potential seeds for microsatellites, or if some STRs in L1s were difficult to identify and include in the reference file. Alternatively, the genomic location of TE insertions may play a large role in the mutation rate of the associated STRs. *Alu* elements, particularly older insertions, have been shown to insert in more GC-rich regions of the genome, while L1 insertions have been found in more AT-rich regions (Brookfield 2001; Sellis et al. 2007). Because GC-rich regions of the genome accrue more mutations (as shown in yeast (Kiktev et al. 2018)), *Alu* elements may contain more STR mutations due to their genomic location rather than something unique about the insertion itself. Additionally, an analysis of poly(A) tail length of *Alu* elements and L1 insertions using Tandem Repeats Finder shows that *Alu* elements (mean = ~21; median = 20) have longer identifiable poly(A) tracts than L1s (mean = ~ 13; median = ~11). These longer poly(A) tails may also contribute to the increased number of STR mutations in *Alu* elements. There are likely many factors influencing the relationship between STRs and TEs, and this relationship should be further investigated.

Previous studies of mutation dynamics have noted a male bias for de novo single nucleotide variants (Kong et al. 2012; Goldmann et al. 2016; Sasani et al. 2019). From our analysis, we identified slightly more male than female transmissions for all *de novo* STR mutations, but this excess was not significant. Among families, we find that more families show male transmission bias than female transmission bias (Figure 5). Comparing the male and female transmissions within a single family shows that either males or females may have statisically significant excess transmissions, but also that these cases are infrequent. This finding is, again, quite similar to the results shown for single nucleotide variants in these families. Due to the challenges and high false positive rate associated with genotyping STRs, we were unable to examine *de novo* mutations in the third generation of individuals within these families. This also prevented us from examining the effect of parental age (within a single family) on mutation rate. Collection of the fourth generation of the CEPH pedigrees is now underway and will allow new analyses to better understand how parental age, genetic background, and differences within DNA repair genes play a role in altering mutation rates.

Comparing the size of the mutation to the motif length of the STR, we find that most of the mutations occur in a stepwise fashion (Figure 7). This mutation pattern has been noted in other studies (Sajantila et al. 1999; Kayser et al. 2000). Regardless of the repeat motif length, we found a rapidly decreasing number of mutations as the step size increased. Slippage has been the proposed mechanism for most single step STR mutations, though larger STR mutations may be caused by other mechanisms and should be further investigated (Fan and Chu 2007). Additionally, we find that most of the identified *de novo* mutations occur in perfect repeats (Table 1). This has been found in previous work and is hypothesized to be due to increased replication slippage in these perfectly repeating regions (Kruglyak et al. 1998; Sun et al. 2012).

We were able to utilize the unique structure of the CEPH pedigrees to better understand the mutational dynamics of STR loci in healthy individuals. As sequencing technology (e.g., long-read sequencing) and computational methods for the detection and genotyping of STRs improve, precisely genotyping longer STRs will improve the estimate of the mutation rate. Future analyses of large pedigrees from diverse populations may uncover additional variation in STR mutation rates, as we have only examined families of European ancestry. Our study provides new perspectives on the dynamics of STR mutations and highlights the need for larger sample sizes and novel tools to investigate this underappreciated portion of the genome.

## Methods

### Sequencing data

Whole-genome Sequencing (WGS) data were available from 599 individuals from 33 families (Sasani et al. 2019). These genomes were sequenced to ~30x coverage, with complete coverage data for each genome used in this study shown in Supplemental Table 3. Coverage data for each file were generated using covstats from the goleft package (https://github.com/brentp/goleft). These data are available with controlled access through dbGaP (phs001872.v1.p1).

### Short Tandem Repeat Genotyping and Basic Filtering

HipSTR (version 0.6.2) (Willems et al. 2017) was used to genotype short tandem repeats (STRs) in the CEPH sequencing data. HipSTR was run on 29 families, though in some cases not all individuals from a family could be successfully run due to a known issue with HipSTR stemming from difficulty extracting certain filtering tags from BAM files. Most pedigrees contain all four grandparents in the first generation (only four pedigrees are missing one or more grandparents). Some families contain multiple offspring (parents) in the second generation, allowing for the analysis of multiple individuals in some of the 29 pedigrees. Splitting these extended pedigrees produced 36 family units used for some analyses. Each CEPH family was run individually using the default stutter model, as recommended in the HipSTR documentation. The GRCh37 STR reference file from https://github.com/HipSTR-Tool/HipSTR-references/ (which includes approximately 1.6 million loci) was used for all analyses. Filtering methods included with HipSTR and DumpSTR from TRTools (version 3.0.0) (Mousavi et al. 2020) were used to filter the genotypes generated by HipSTR. Specifically, we filtered these genotypes for loci with a minimum quality score > 0.9, maximum flanking indels < 0.15, maximum call stutter < 0.15, minimum call depth of 10, maximum call depth of 1000. All loci overlapping segmental duplications in the genome were removed. With these filtering criteria, we retained loci that were confidently genotyped, did not contain too many flanking indels, and showed little evidence of stutter artifacts in the repeat.

### Validation of Genotypes

STR genotypes were previously generated in the 1990s for 360 markers for individuals in the CEPH pedigrees as part of the Human Genome Project. STR loci were amplified by PCR, and STR genotypes were detected and visualized using an Automated Hybridization and Imaging Instrument (AHII) by modifying the methodology previously used to generate high throughput *de novo* sequence data (Cherry et al. 1994). These loci were identified in hg19 using the UCSC Genome Browser; then the genotypes coded by HipSTR were compared to the genotypes previously coded using AHII. A list of the loci and genotypes generated by AHII and HipSTR are shown in Supplemental Table 4. In total, we analyzed 5 loci in multiple families, allowing for the comparison of 175 genotype calls between the two methods.

### Filtering for *de novo* Mutations

After filtering the genotypes generated with HipSTR, the genotypes for individuals in the second generation of the CEPH families were compared to those of their parents (generation 1) to identify potential *de novo* mutations. For a mutation to be considered for further analysis, we required that at least 10 reads spanned the *de novo* allele in the individual in the second generation. This decreased the number of loci that could be considered, but it provides greater confidence in the called genotype.

Next, each potential *de novo* STR that met these criteria in the second generation was compared to the third generation to ensure that the *de novo* allele was transmitted to multiple individuals (at least two) in the third generation (Willems et al. 2017). Given the high mutation rate of STR loci, requiring that at least two individuals inherited the *de novo* allele increased our confidence in the successful transmission of the *de novo* allele. Further, to ensure the *de novo* allele was transmitted from the individual in which it was identified, we removed from consideration all *de novo* STRs that were shared with the other parent (similar methodology was also used in (Willems et al. 2017)). To calculate the STR mutation rate, we divided the number of *de novo* mutations by the total number of STRs that passed our filters for each trio.

### Identifying perfect and imperfect repeats

To identify which STR loci were perfect and which were imperfect, we used Tandem Repeats Finder (Benson 1999) (Version 4.09.1). BEDTOOLS (Quinlan and Hall 2010) was used to get sequence data for each of the approximately 1.6 million STR loci included in the HipSTR reference file. The sequence data were analyzed with Tandem Repeats Finder with a minimum score requirement of 15 to ensure that even the short repeats included in the HipSTR reference file could be accurately identified. Each motif size (mononucleotide - hexanucleotide) was run separately to be sure we were identifying the correct repeat. Those repeats that had a perfect score for “Percent Matches” in the output file were considered to be perfect repeats. All other repeats were considered to be imperfect. We then used BEDTOOLS to intersect the loci in which we identified *de novo* STR mutations with the location of the perfect and imperfect repeats.

### Genomic Location of *de novo* mutations

After identifying *de novo* mutations in STRs, we intersected the location of these mutations with exons, introns, 5’-UTRs, and 3’-UTRs using BEDTOOLS (Quinlan and Hall 2010). We used the UCSC Genome Browser to find the location of genes in hg19. The different regions of genes, as listed above, were run separately to determine if the *de novo* STRs intersected with any component of a gene. A similar procedure was used to find the location of active transposable elements (*Alu* elements, L1, and SVA) in hg19. These loci were intersected with the identified *de novo* mutations to determine the frequency with which transposable elements were the sites of these mutations.

### Identification of Poly(A) tails in *Alu* elements and L1s

To determine the length of identifiable poly(A) tails in *Alu* elements and L1s, we obtained fasta files of the sequences for each insertion (with an additional 40 bp of flanking sequence on the 3’ end) in hg19 from the UCSC Genome Browser. Short tandem repeats in each element were identified with Tandem Repeats Finder. The results were then filtered to only include those that were found near the end of the insertion and had a repeat motif of “A”. From the filtered results, the mean and median length of the identifiable poly(A) tails were determined.

### STRdiff

STRdiff (https://github.com/ScottWatkins/STRdiff.jl) was used to evaluate the characteristics of *de novo* STR mutations found in parents of the CEPH families. This program leverages the three-generation structure of the CEPH pedigrees to infer the sex and haplotype of the grandparent transmitting the *de novo* STR allele and subsequent size change, in base pairs, between the original and the *de novo* alleles.

Input to STRdiff is a variant call format (vcf) file containing two sets of grandparents, two parents, and all offspring. A region surrounding the *de novo* STR is first extracted and then phased in all possible trios in the family using the Beagle software package (Browning and Browning 2007). Because the de novo STR is a mutational event that creates a misinherited allele, the *de novo* allele cannot be phased directly. Instead, the haplotype(s) carrying the novel STR allele are first identified in multiple offspring. A consensus haplotype is created from all offspring haplotypes that carry the *de novo* allele. Using a consensus haplotype helps to reduce mismatches caused by rare alleles, sequencing/genotyping errors, and inferred recombination events. The consensus haplotype is then compared to each of the four phased grandparental chromosomes of the parent that harbors the *de novo* mutation. The fraction of alleles contained on each grandparental haplotype that match the offspring consensus haplotype bearing the de novo STR allele is calculated. Match probabilities are used to identify the most likely grandparental chromosome with the original STR allele that produced the mutational event.

Depending on the chromosomal location and family, regional haplotypes may be highly similar. To improve the number of *de novo* STRs for which a transmitting grandparental haplotype could be reliably identified, STRdiff was run over a range of haplotype sizes (10, 20, …, 300kb). By using this broad range of haplotypes, a probable haplotype solution was found for 2361 of 2456 (96%) de novo STRs. Of these 2361 haplotypes, 2202 (93%) were uniquely resolved from other haplotypes at a minimum of ≥ 10% of all polymorphic sites found along the length of the haplotype. Additionally, we tested the concordance between haplotypes constructed directly from WGS sequence and those based on 1.1M SNPs common Illumina array SNPs extracted from the WGS sequence. There was >94% concordance for sex assignment for the transmitting grandparent. Assignments that differed were most often due to a high similarity among parental and grandparental haplotypes at that locus and minor phasing differences among the inferred haplotypes.

To obtain a more rigourous independent estimate of the accuracy of STRdiff, a subset of 24 *de novo* transmission events were examined in IGV (Robinson et al. 2011). These loci, along with details of the STRdiff prediction for each, are shown in Supplemental Table 5. The examined loci were selected randomly, but we required that these loci had sufficient read depth, low stutter at the STR locus, and usually a nearby SNP to allow for independent read-based phasing. Mononucleotide repeats were particularly difficult to analyze through IGV. The STRdiff and IGV read-based phasing predictions for the parent transmitting the de novo STR allele matched in 23/24 (>95%) of the examined loci. The allele size changes, when predicted by STRdiff, was correct for 19/21 (~90%) loci. The IGV images associated with each locus are shown in Supplemental Figure 4.

### Size change of STR mutations

The stepwise change for each class of STR repeat (mono, di, …) was calculated in STRdiff using information from the vcf file. The sequence for the transmitting grandparent’s allele and the *de novo* allele in the offspring was obtained directly from the vcf file. The absolute value of the difference in base pairs between these alleles was divided by the allele class size to determine the number of steps. For example, a dinucleotide repeat allele, CACACA, that becomes a CACACACA repeat allele is scored as a one repeat change, |(6-8)/2| = 1. Repeat size changes could not always be resolved due to similarity in haplotypes and STR homozygosity in the grandparents, parents, or offspring. In total, repeat size changes were determined for 1533 mutational events.

## Acknowledgments

This work was supported by NIH R35GM118335 (to LBJ), NIH/NRSA T32HG008962 (to CJS), and NIH K99HG011657 (to CJS). We also thank the Utah Genome Project, the George S. and Dolores Doré Eccles Foundation, and the H.A. and Edna Benning Society for sequencing funds. We thank all of the Utah individuals who participated in the CEPH consortium. Additionally, we would like to thank members of the Jorde and Quinlan labs for their helpful feedback and discussion on this project.

## Confilcts

The authors have no conflicts to disclose.

---

**Supplemental Figure 1.**
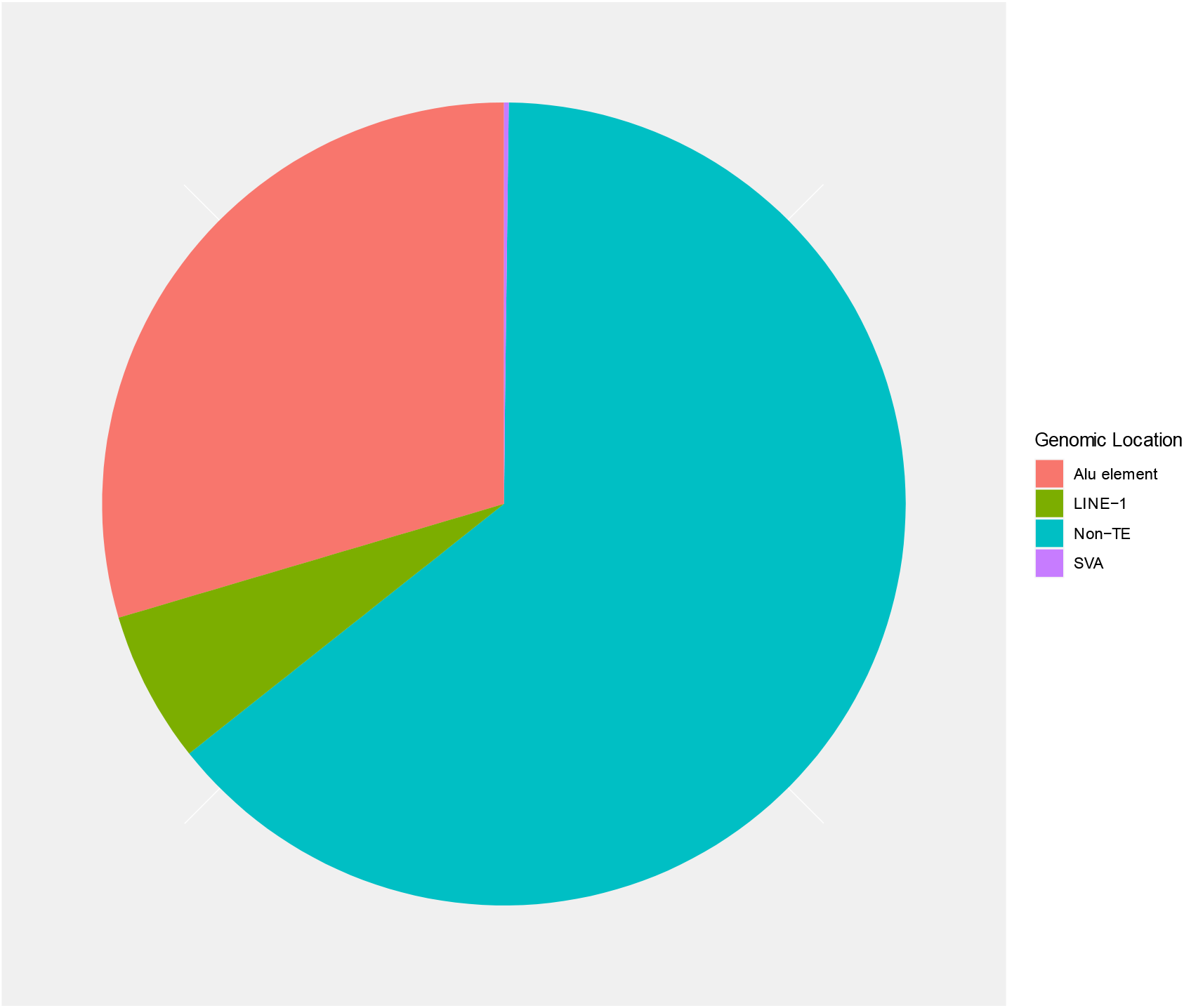
*De novo* STRs that intersect transposable elements. *Alu* elements intersect approximately 30% of the *de novo* STRs that were identified in this study (red). LINE-1 (L1) insertions show a smaller proportion (green), with those that intersect SVA insertions shown in the small purple sliver. The remaining portion of the chart (blue) shows the *de novo* STRs that intersect non-transposable element (TE) regions of the genome.

**Supplemental Figure 2.**
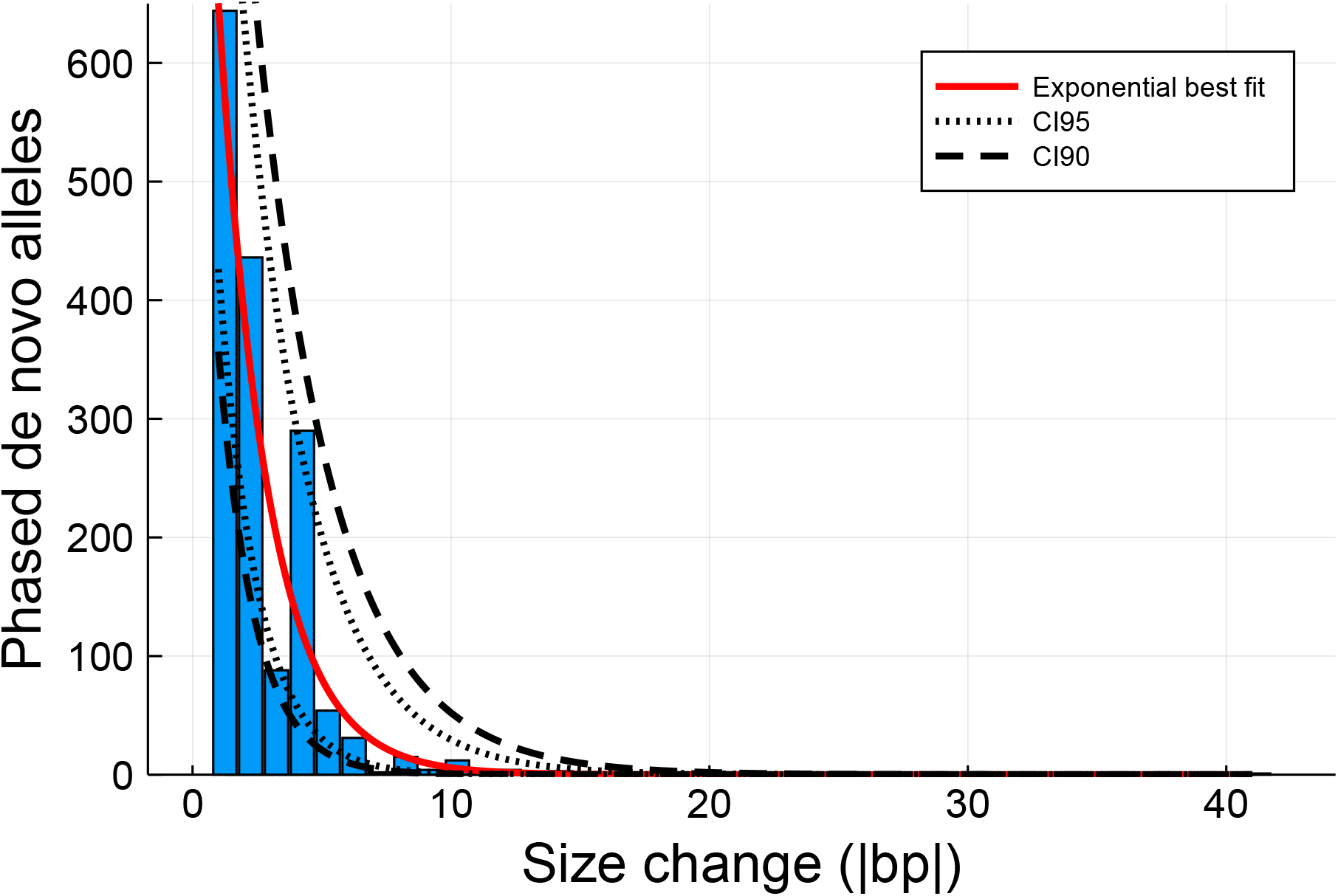
Stepwise changes observed all *de novo* STR alleles follow a negative distribution. All motif lengths except for trinucleotide repeats fall within the 95 percent confidence interval. Trinucleotide repeats fall within the 90 percent confidence interval.

**Supplemental Figure 3.**
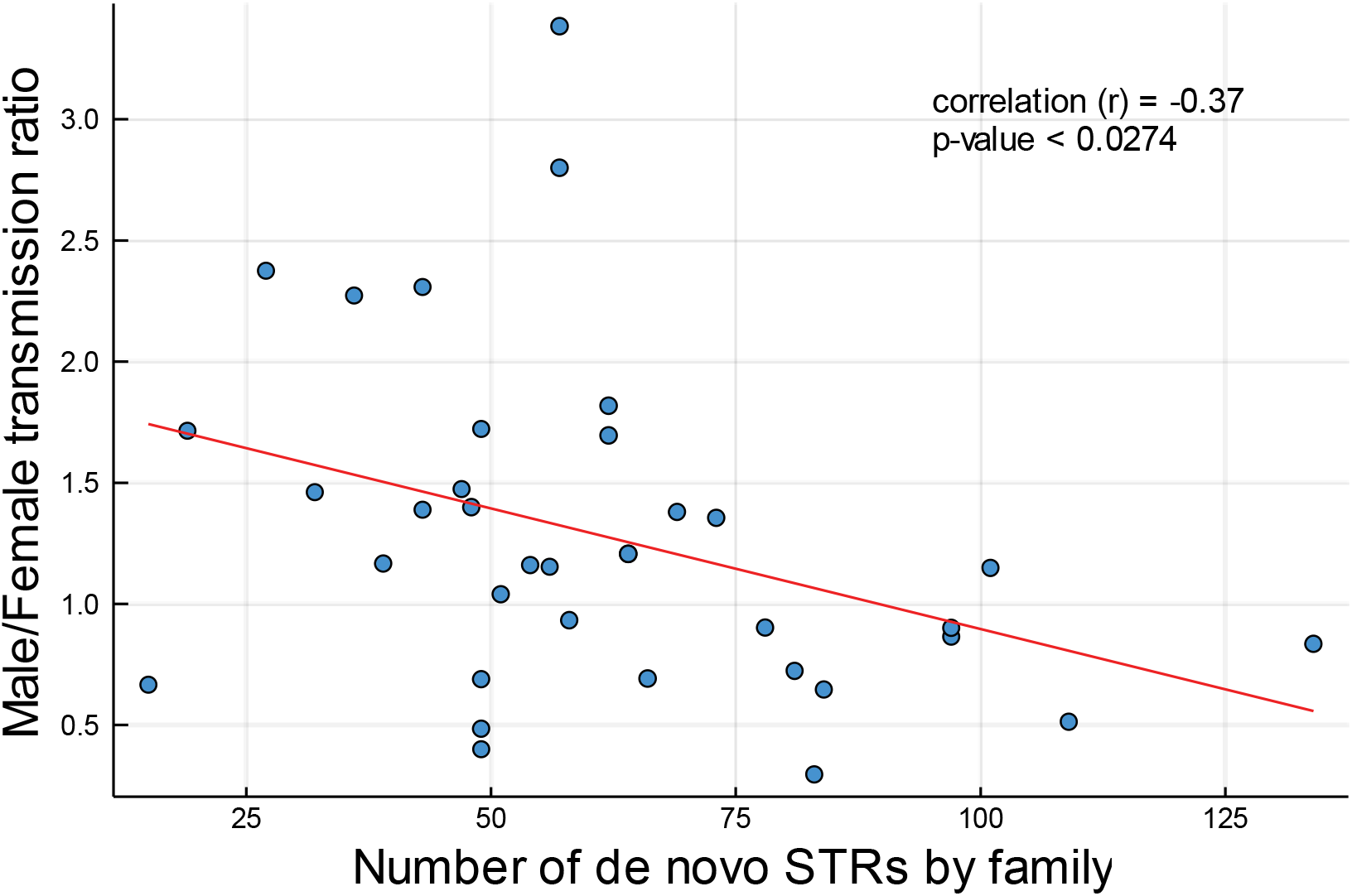
Observed male/female ratio decreases with increasing number of de novo STRs. For 36 CEPH families, the ratio of de novo mutations transmitted by males compared to females decreases with an increasing number of de novo STRs discovered in the family.

**Supplemental Table 1.** Distribution of de novo STRs by genomic location.

|  | Observed | Expected | Odds Ratio<br>(CI 95%) | P-value |
| --- | --- | --- | --- | --- |
| **Coding exons** | 2 | 31 | 0.06<br>(0.01 - 0.25) | < 1e-6* |
| **5'-UTR** | 31 | 36 | 0.86<br>(0.51 - 1.43) | > 0.623 |
| **3'-UTR** | 15 | 17 | 0.88<br>(0.41 - 1.88) | > 0.860 |
| **Introns** | 1233 | 1119 | 1.18<br>(1.06 - 1.32) | < 0.002* |
| **Intergenic** | 1466 | 1544 | 0.89<br>(0.80 - 0.99) | > 0.037 |

(\* significant with Bonferroni correction; Fisher's exact test)

**Supplemental Table 2.**
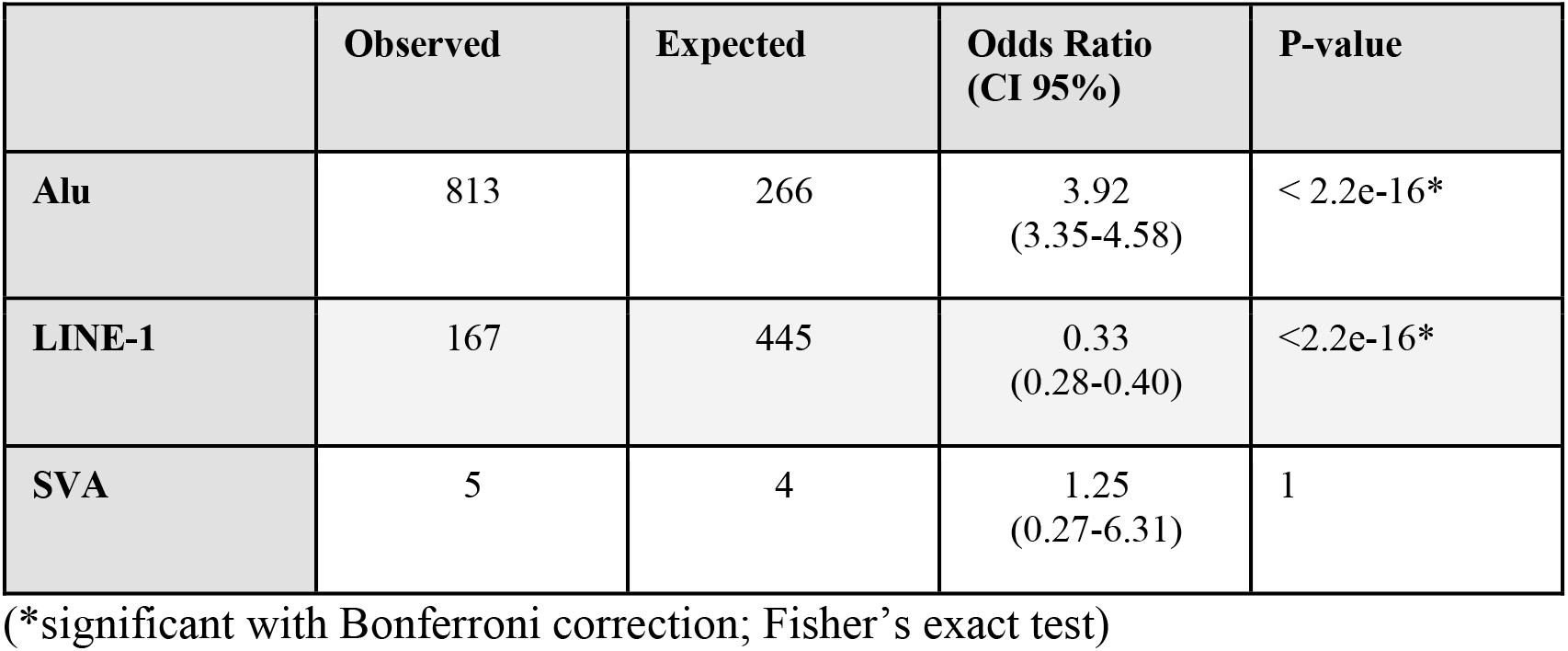
Distribution of *de novo* STRs in transposable elements.

**Supplemental Table 5.**
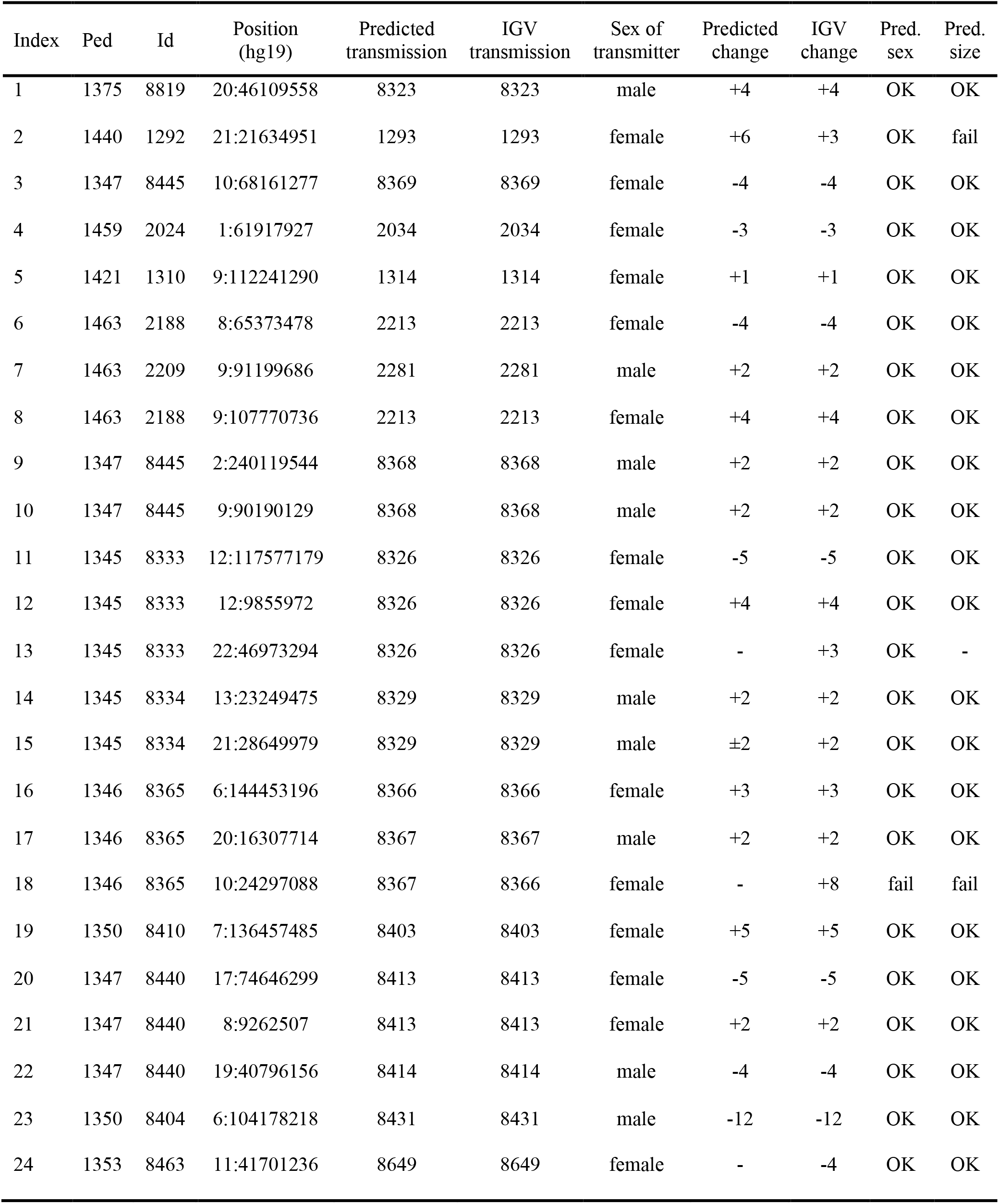
Read-based estimates of STRdiff accuracy using IGV.

